# Targeting Metabolic Dependencies in Acute Myeloid Leukemia: A Dual Strategy using Arsenic Trioxide and Artesunate

**DOI:** 10.1101/2024.06.23.600168

**Authors:** Nithya Balasundaram, Arvind Venkatraman, Yolanda Augustin, Hamenth Kumar Palani, Clement Regnault, Anu Korula, Uday Prakash Kulkarni, Eunice Sindhuvi Edison, Poonkuzhali Balasubramanian, Biju George, Aby Abraham, Sanjeev Krishna, Vikram Mathews

**Affiliations:** Department of Haematology, Christian Medical College, Vellore, India; Clinical Academic Group in Institute for Infection & Immunity, St George’s University of London, London, United Kingdom; St George’s University Hospitals NHS Foundation Trust, United Kingdom; Institut für Tropenmedizin, Universitätsklinikum Tübingen, Tübingen, Germany; Centre de Recherches Médicales en Lambaréné (CERMEL), Lambaréné, Gabon; MVLS Shared Research Facilities, University of Glasgow, Glasgow, United Kingdom

**Keywords:** acute myeloid leukemia, artesunate, arsenic trioxide, mitochondria, uncoupler, iron, calcium, acute myeloid leukemia

## Abstract

Acute myeloid leukemia (AML) remains a difficult disease to cure despite recent advances. Off-target side effects of therapy, especially prolonged cytopenia, lead to significant morbidity and mortality. There is an increased recognition that in AML, there is a potentially vulnerable dependence on OXPHOS metabolism, more so in the leukemia stem cell compartment (AML-LSC) that could be exploited. Drug re-purposing screens have suggested the possible role of artesunate (ART) in inhibiting mitochondrial respiration and arsenic trioxide (ATO) in inhibiting glycolysis. Here, we explore the potential of using a combination of ART and ATO to treat AML. Through in-vitro and in-vivo studies, we demonstrate the potential of this combination along with hypo-methylating agents and other mitocans, such as BCL-2 inhibitor venetoclax, to be used in the treatment of AML with minimal off-target effect on normal hematopoietic stem cells (HSC). These observations warrant further exploration of such novel combinations in clinical trials.

## Introduction

Acute myeloid leukemia (AML) is the most common leukemia in adults. It has a 5-year survival rate of 25-40%, with little improvement in the past 30 years^1^. Treatment usually involves an intensive cytoreductive 7+3 induction therapy combining a nucleoside analogue for seven days (cytarabine) with an anthracycline for 3 days (daunorubicin, idarubicin, or doxorubicin) or the more recently introduced low-intensity combination of BCL-2 inhibitor (venetoclax (VEN)) and a DNA methyl transferase inhibitor (5-azacytidine or decitabine)^2,3^. AML is an oligoclonal disease, genetically and morphologically heterogenous, which allows escape from targeted therapies such as FLT-3 inhibitors (Midostaurin and Gliternib) and IDH1/2 inhibitors (Enasidenib and Ivosidenib)^4^. This emphasizes the need to optimize combination therapies and treatment strategies to cure this disease.

Otto Warburg first observed that cancers share the metabolic characteristic of increased glycolysis without hypoxia^5,6^. Although he attributed increased glycolysis to mitochondrial dysfunction, increased glycolysis is anaplerotic by replenishing substrates for synthesizing nucleosides, lipids, and proteins. In contrast, in AML, it is recognized that there is an increased dependence on mitochondrial respiration, especially in the AML leukemia stem cell compartment^7,8^. However, targeting mitochondrial dysregulation using relatively specific inhibitors of complex-I (IACS-010759) and a DHODH inhibitor (beraquinar) and other less specific interventions (metformin and tigecycline) has not translated from experimental models to therapeutic benefits in patients ^9–11^. Partly, this is due to the metabolic adaptability of AML cells, particularly therapy resistant cells, as previously documented^12^.

These observations suggest that targeting the electron transport chain (ETC) is insufficient to lethally impact the central energy metabolism (CEM) of AML cells, particularly those resistant to conventional treatments. Defining CEM pathophysiology more comprehensively in AML may improve treatment options by suggesting rationally derived combination therapies targeting therapy-resistant cells. For example, combining FCCP with glycolytic inhibitors such as arsenic trioxide (ATO) or 2-Deoxy-D-glucose (2-DG) promoted apoptosis in otherwise ATO-resistant acute promyelocytic leukemia (APL) cells^12^. We demonstrated a similar effect in AML cell lines and primary AML cells that were inherently resistant to ATO^12^. The challenge with the combination of ATO+FCCP was the lack of selective specificity to differentiate normal hematopoietic stem cells (HSC) and normal peripheral blood mononuclear cells (PBMNC) from the leukemic cells ^12^.

We hypothesized that combining ATO, and cancer cell specific ETC inhibitors may result in lethal CEM when individual pathway inhibitions are ineffective. Based on existing drug repurposing screens that identify shifts in metabolism from mitochondrial respiration to glycolysis we evaluated artemisinin as a potential alternative for FCCP to selectively target the leukemic cells over their normal counterparts^13^. CEM is not only regulated by flux and availability of metabolites, but it is also highly regulated by the co-factors such as iron and calcium that are often increased in cancer cells in parallel with metabolic derangements^14–17^.

We hypothesized that exploiting these alterations would have the potential to selectively target cancer cells. We therefore examined the combination of ATO and artesunate (ART) on AML cells using multidisciplinary approaches. ART is an antimalarial agent that is being increasingly repurposed for its potential as an anti-cancer therapy. Our research findings are driving an ongoing clinical development program in this promising area.

## Results

### ATO and ART selectively disrupt the CEM of AML cells

ATO is known for its activity on PML-RARA oncoprotein, and its anti-leukemic activity in APL. However, we found that ATO inhibits glycolysis, and that non-APL AML cells utilize OXPHOS, circumventing ATO-mediated cell death. The efficacy of ATO on AML cells is highly correlated to the competitive glucose analogue 2-deoxy glucose (2-DG) (figure 1a). As we observed metabolic adaptability in ATO- resistant AML cells, we hypothesized that a two-pronged approach incorporating mitochondrial OXPHOS uncouplers or inhibitors combined with ATO would be required to eliminate the leukemic cells. We evaluated the combination of ATO (2uM) with different peroxidic antimalarials (5uM) and anti-metabolites (5uM) based on the published literature^13^. ART synergized with ATO and elicited a profound anti-leukemic effect on U937 cells (Fig1b). This synergistic combination-maintained efficacy on AML, APL, ALL cell lines, and their subtypes, with minimal perturbations on normal HSC and PBMNC (figure 1c). Giemsa stained U937 cells treated with ATO or ART did not show any differentiation (Figure S1a). We assessed if adding ATO and ART as sequential single agents would further enhance their efficacy compared to concurrent treatment with ATO+ART. Concurrent treatment was superior to sequential single agents of ATO, followed by ART or vice versa (figure 1d).

**Figure 1:**
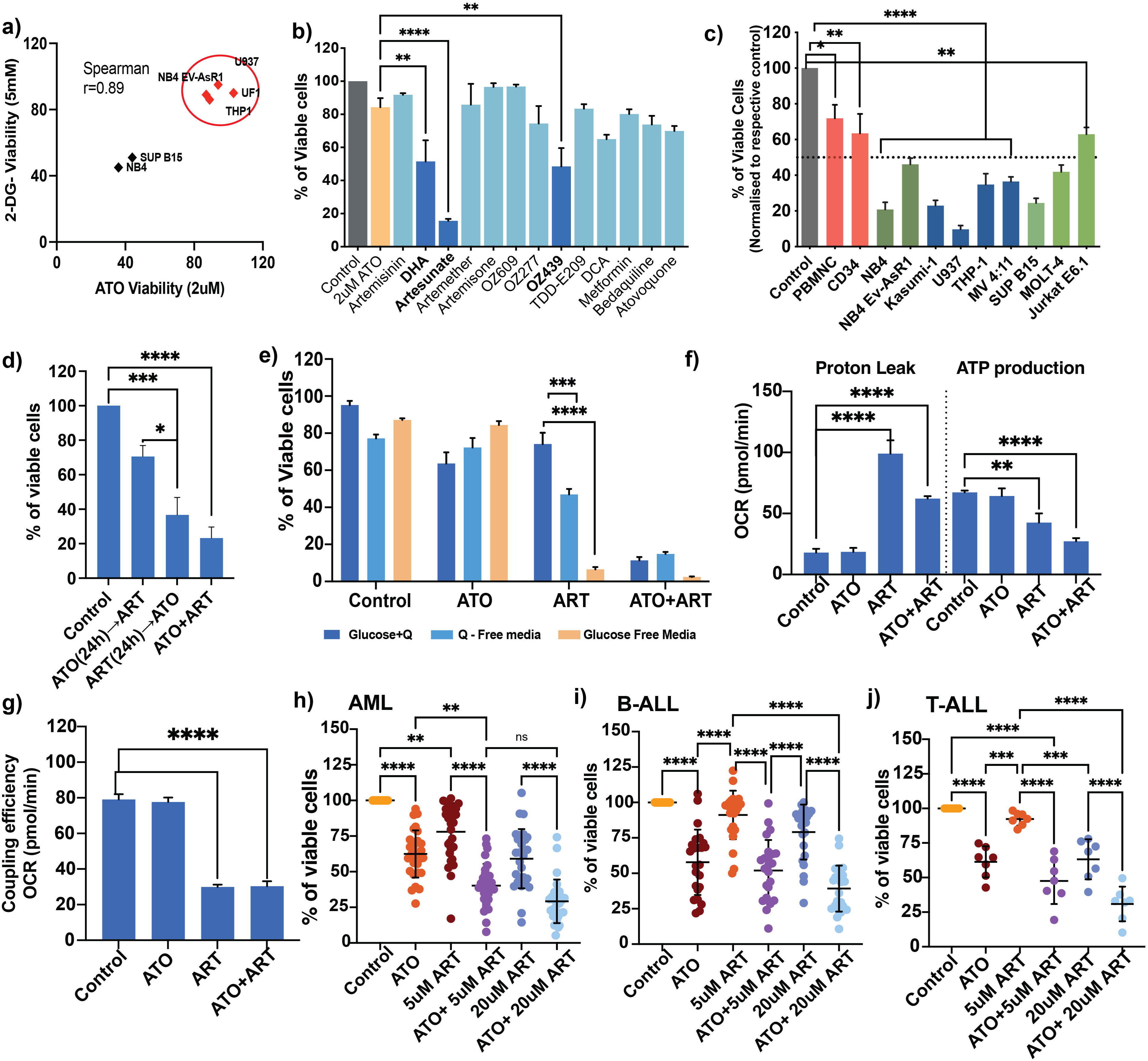
ATO and ART disrupt the CEM of AML cells with minimal off-target effects on normal cells: a) Spearman correlation of AML cell lines viability post-2-DG (5mM) and ATO (2uM) treatment for 48h (n=3). b) Viability of U937 cells treated with different peroxidic antimalarials and anti-metabolites in combination with ATO (Light and dark blue represent the combination of ATO, and a yellow bar represents single agent ATO) (n=3; ATO=2uM; antimalarials=5uM; DCA=10mM; Metformin= 5mM; Bedaquiline and Atovaquone = 10uM). c) Viability of APL (Dark green: NB4, NB4 EV-AsR1), AML cell lines (Dark blue: U937, THP-1, MV411, Kasumi-1), ALL (Light green: SUP B-15, MOLT-4 and Jurkat e6.1) and normal PBMNC and CD34 positive cells (Red; p<0.05) treated with a combination of ATO and ART for 48h. Results were normalised to their respective control. For values <50 % (dotted line), p <0.0001; (normal cells n=3; cell lines n=7 myeloid; n=3 for lymphoid ATO =2uM; ART = 5uM). d) Viability of U937 cells with sequential (ATO or ART as single agents for 24h followed by ART or ATO) and combination treatment of ATO and ART for 48h (n=3; ATO = 2uM; ART= 5uM). e) Viability of U937 cells to ATO and ART in the presence and absence of glucose and glutamine(Q) (n=3; ATO = 2uM; ART= 5uM). (f -g) Proton leak, ATP production (f) and coupling efficiency (g) of U937 cells treated with ATO and ART followed by seahorse mito stress analysis (n=3; ATO = 2uM; ART= 5uM). (h-j) Viability after ATO and ART treatment for 48h; Primary AML (h), B-ALL (i) and T-ALL (j) cells AML (n=28), B-ALL (n=24) and T-ALL (n=7). ATO was used at 2uM, and ART was at 5uM and 20uM. Data are presented as mean ± SEM. n.s., P > .05; ∗P < .05; ∗∗∗P < .001; ∗∗∗P < .0001, with a two-tailed unpaired t-test or one-way analysis of variance.

To strengthen our dual approach targeting the leukemic cells, we evaluated the effects of ATO and ART on AML cells cultured in glutamine and glucose-free conditions. As hypothesized, ART as a single agent had a profound impact in the glucose-free condition, equivalent to the combination of ATO and ART in the presence of glucose and glutamine (figure 1e). We also assessed the combination of GLS (glutaminase inhibitor- CB839) and etomoxir (Fatty acid oxidation inhibitor) with ATO or ART. There was no synergistic effect of these agents with ATO or ART (Figure S1d). However, we observed that ART increased GLUT- 1, GLS, and CD36 expression, whereas ATO decreased GLUT-1 and did not change CD36 and GLS expression (Figure S1e, f and g). Mito stress analysis of U937 cells treated with ATO and ART revealed that ART increased proton leak and decreased ATP production (figure 1f) and coupling efficiency (figure 1g). In contrast, ATO alone did not affect these parameters (figure 1f and g). Reactive oxygen species levels were not elevated by treatment with ATO and ART at 6h (Figure S1b, however by 24h there was a reduction in the ROS levels after ATO, ART, and ATO+ART compared to control U937 cells (Figure S1c).

We then validated the therapeutic effect of ATO+ART on primary bone marrow samples of AML, BCP-ALL and T-ALL obtained at diagnosis. ATO+ART had dose-dependent anti-leukemic activity across all cell types of acute leukemia (figure 1h, 1i and 1j).

### Cellular targets of artesunate in AML

To identify the cellular targets of ART, we employed chemical drug proteomics to label ART-interacting proteins. U937 cells (myeloid lineage, pro- monocytic) were treated with biotinylated ART (having demonstrated similar activity to the parent compound) for 6 hours, and the ART targets were affinity purified by streptavidin beads and identified with LC/MS (figure 2a). Gene ontology analysis for cellular component enrichment showed that ART interacts with many cellular components (figure 2b). Pathway analysis of the enriched proteins revealed that they are involved in c-myc, OXPHOS, unfolded protein response and fatty acid metabolism (figure 2c, Supplementary table 1). We assessed the metabolic perturbation caused by ART by performing an untargeted metabolomics on the U937 cells treated with ART for 24 hours, we noted that acylcarnitines, triglycerides, lysophosphotidylcholine, ceramide-related metabolites are dysregulated in comparison to control (figure S2, supplementary table 2). To validate our findings from chemical drug proteomic and metabolomics, we performed knockdown (KD) of Very Long Chain acyl-CoA Dehydrogenase (VLCAD; ACADVL) that has been reported to be overexpressed and a druggable target in AML^18^. In U937 cells, lentiviral short hairpin RNA (shRNA) was used to knock down VLCAD expression (figure 2d). VLCAD knockdown has been shown to increase glycolysis in AML cells, supporting that we observed sensitivity of knockdown cells to ATO as a single agent (figure 2e). Electron microscopy revealed disrupted mitochondrial structures and an increase in lipid droplets in these cells compared to cells transduced with scrambled shRNA (figure 2f).

**Figure 2:**
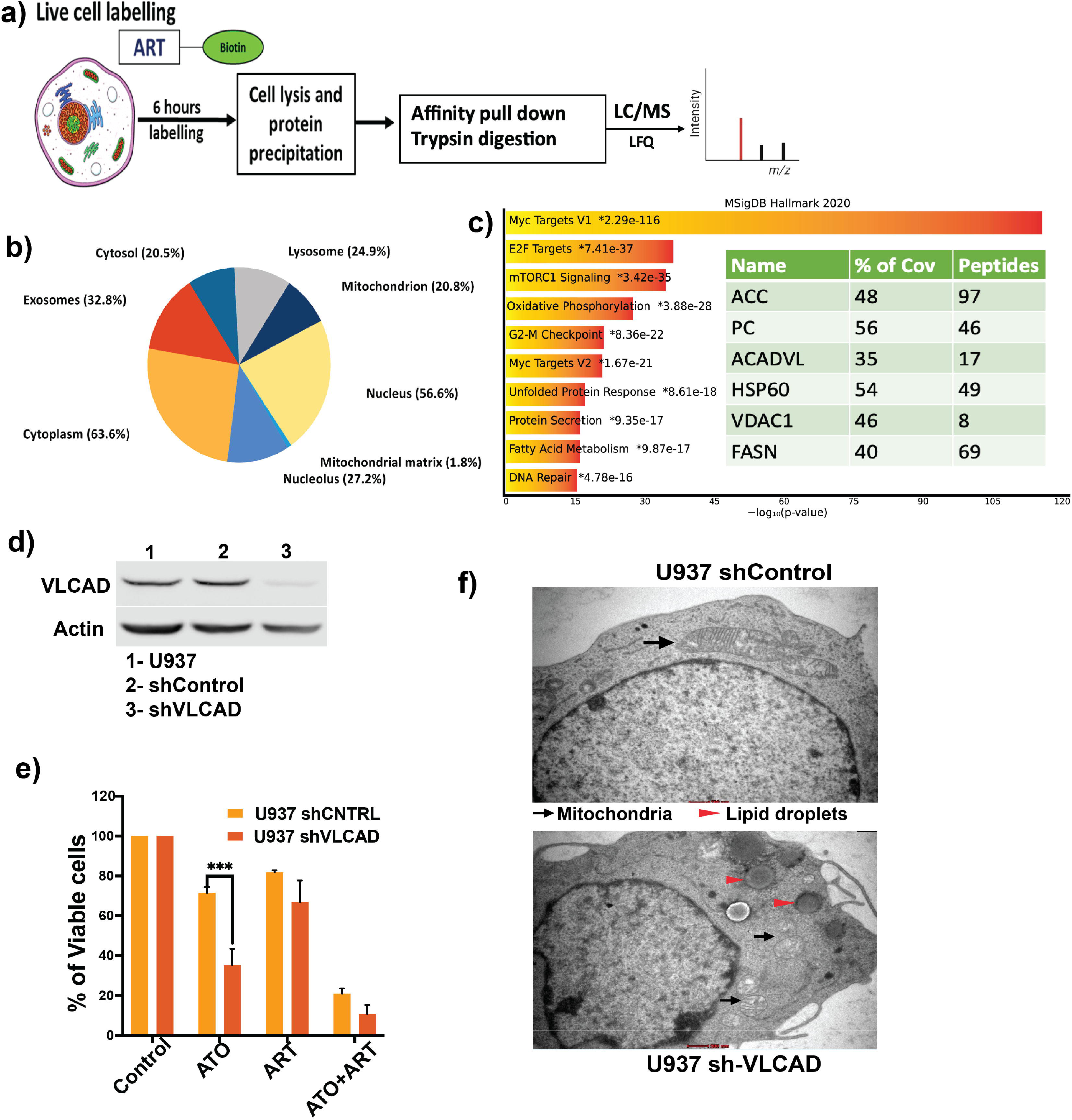
Cellular targets of artesunate in AML cells: a) General workflow of the chemical proteomics approach where the leukemic cells are treated with biotinylated ART (50uM, 6hrs) and streptavidin pull-down followed by mass spectrometry. b) Pie chart illustrating the cellular component enrichment from gene ontology (GO) analysis of identified direct protein targets. c) KEGG pathway enrichment analysis of metabolites downregulated by ART in the U937 cells. (x-axis; enrichment ratio of the KEGG pathway and the intensity of red indicates the p-value highest to lowest). Pathway analysis of the protein targets and a table of proteins with the highest overall coverage. ACC, acetyl-CoA carboxylase 1; PC, pyruvate carboxylase; ACADVL, very long-chain specific acyl-CoA dehydrogenase; HSP60, heat shock protein 60 kDa; VDAC1, voltage-dependent anion-selective channel protein 1; FASN, Fatty acid synthase N. % Cov, per cent protein sequence coverage with the identified peptides. d) Immunoblot of VLCAD in U937 cells transduced with scrambled control shRNA and shRNA against VLCAD. e) Representative electron micrographs of shVLCAD and shCNTRL U937 cells. Scale bars represent 0.5um. f) Viability of U937 shCNTRL and shVLCAD cells treated with ATO and ART for 48h (cultured in complete media) (n=4; ATO =2uM; ART=5uM) Data are presented as mean ± SEM (n = 3). n.s., P > .05; ∗P < .05; ∗∗∗P < .001, ∗∗∗P < .0001, vs sh CNTRL with a two-tailed unpaired t-test or one-way analysis of variance.

### Mitophagy diminishes ART sensitivity

ART-treated U937 cells showed disruption of mitochondrial structures with prominent loss of cristae, and fragmentation visualized under the electron microscope (figure 3a) and confocal microscopy (figure 3b). The morphological changes of the mitochondria suggest that the damaged mitochondria are probably actively cleared in response to ART-induced stress via mitophagy; hence, adding a mitochondrial fission inhibitor mdivi-1 (mitochondrial division inhibitor 1) in combination with ART should enhance the anti-leukemic properties of ART. As expected, mdivi-1 synergized with ART and enhanced cytotoxic effects on U937 cells and primary bone marrow AML specimens (figure 3c, and 3d).

**Figure 3:**
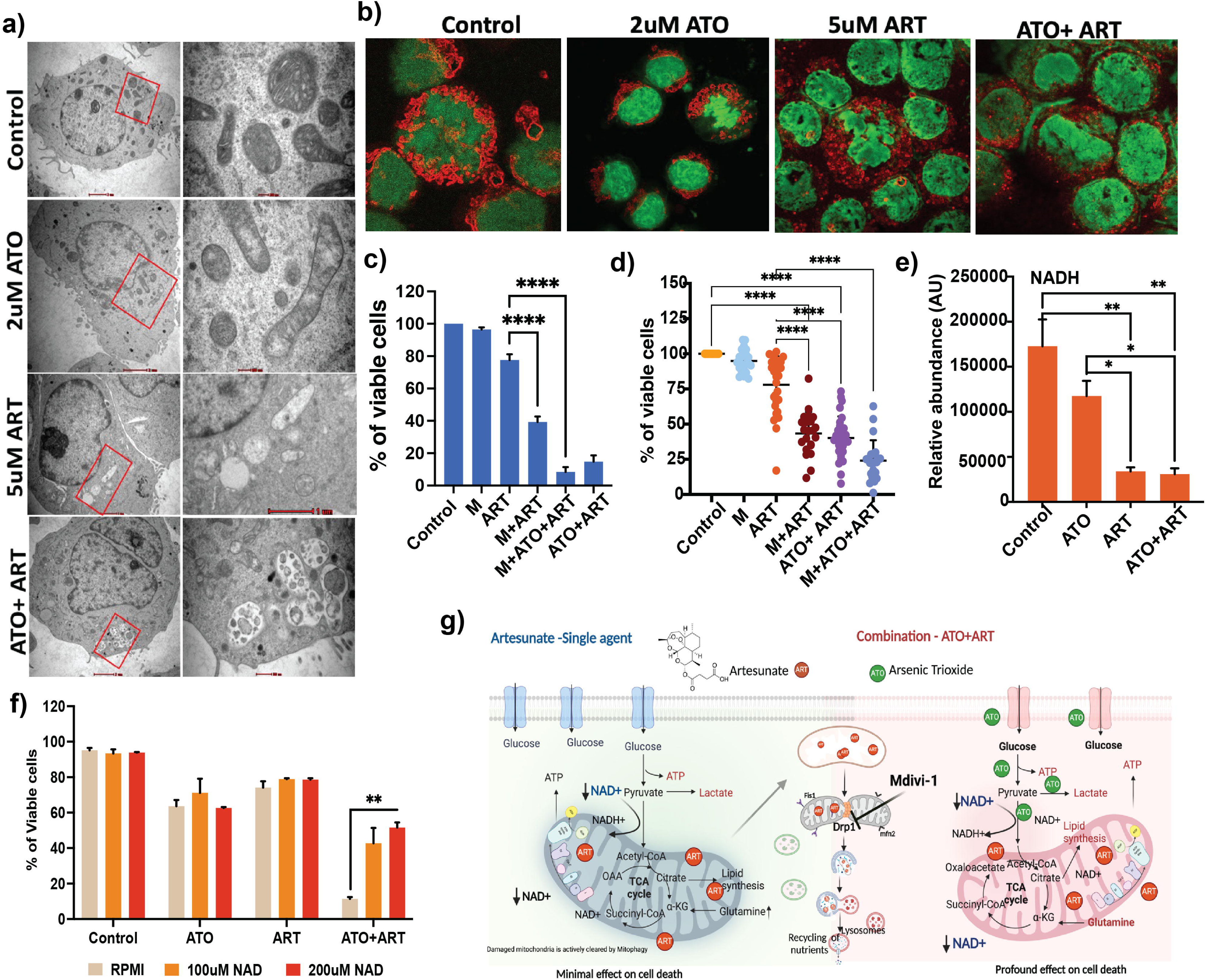
ART disrupts the mitochondrial dynamics and architecture of AML cells: a) Representative electron micrographs of U937 treated with 2uM ATO, 5uM and ATO+ART or control for 24h. Scale bars represent 2μm. b) Representative confocal microscopic images of mitotracker red CMxROS U937 cells treated with ATO, ART and ATO+ART or control for 24h. Stained cells were visualized by FV3000 Inverted Confocal Laser Scanning Microscope, 63× oil immersion objective) c) Viability of U937 cells treated with mitochondrial fission inhibitor (mdivi-1), ART, ATO, and combination for 48h (n=5; ATO =1uM; ART =5uM; mdivi-1 = 10uM). d) Viability of primary AML cells treated with mitochondrial fission inhibitor (mdivi-1), ART, ATO and combination for 48h (n=22; ATO =2uM; ART =5uM; M = 10uM). (e) LC-MS analysis of NADH levels in U937 cells treated with ATO, ART and ATO+ART for 24h (n=3; ATO=2uM; ART=5uM). f) Viability of U937 cells treated with ATO, ART and ATO+ART when supplemented with NAD^+^ for 48h (n=3; ATO=2uM; ART= 5uM; NAD^+^ = 100uM and 200uM) g) Model illustrating the observed effects of the combination ATO+ART: ART accumulated and promoted prominent structural changes in mitochondria. ART uncoupled OXPHOS from glycolysis and reduced cellular NAD+, with minimal effect on leukemic cell survival. However, ART in combination with ATO (glycolytic inhibitor) or mdivi-1(M – Drp1 – Mitophagy inhibitor) or in glucose-deprived conditions promoted cell death via metabolic catastrophe (created with BioRender.com) Data are presented as mean ± SEM. n.s., P > .05; ∗P < .05; ∗∗∗P < .001; ∗∗∗P < .0001, with a two-tailed unpaired t-test or one-way analysis of variance.

NAD^+^ pool is critical for CEM by enabling the cells to adapt to nutrient perturbations and maintain redox states. In our metabolomics data, we noted that NADH levels were reduced by the treatment of ART (figure 3e). Further, we assessed whether increasing the intracellular NAD^+^ would reduce the efficacy of ATO+ART. As expected, we observed that supplementation of NAD^+^ partially reduced the efficacy of the combination in U937 cells (figure 3f). Collectively, these results demonstrate ART promotes uncoupled respiration and mitochondrial damage and in combination with ATO, which inhibits compensatory glycolysis, promotes cell death, as illustrated in the figure 3g.

### Intracellular iron determines the specificity of ATO+ART

Iron plays a vital role in the antimalarial activity of artemisinins by catalysing activation of the endoperoxide moiety. It is recognized in leukaemias that the leukemic cells have enhanced uptake of iron via transferrin receptor(TFRC), decreased efflux and increased cellular content of iron for cell proliferation^19^. It has also been noted that TFRC expression in cancer cells correlates with susceptibility to ART^20^. Deferoxamine (DFO), an iron chelator, diminished the antimalarial activity of artemisinins by chelating the intracellular iron required for their activity ^21–23^. Hence, we used iron chelators selective for different intracellular compartments such as deferiprone (DFP; mitochondria and cytosol), DFO (lysosomes) and 2’2’ Bipyridyl (BIP; cytosol) with ATO + ART to assess the importance of subcellular compartments of iron for anti-leukemic activity. DFO, significantly abrogated the anti-leukemic activity of ATO+ART, whereas DFP and BIP did not (figure 4a).

**Figure 4:**
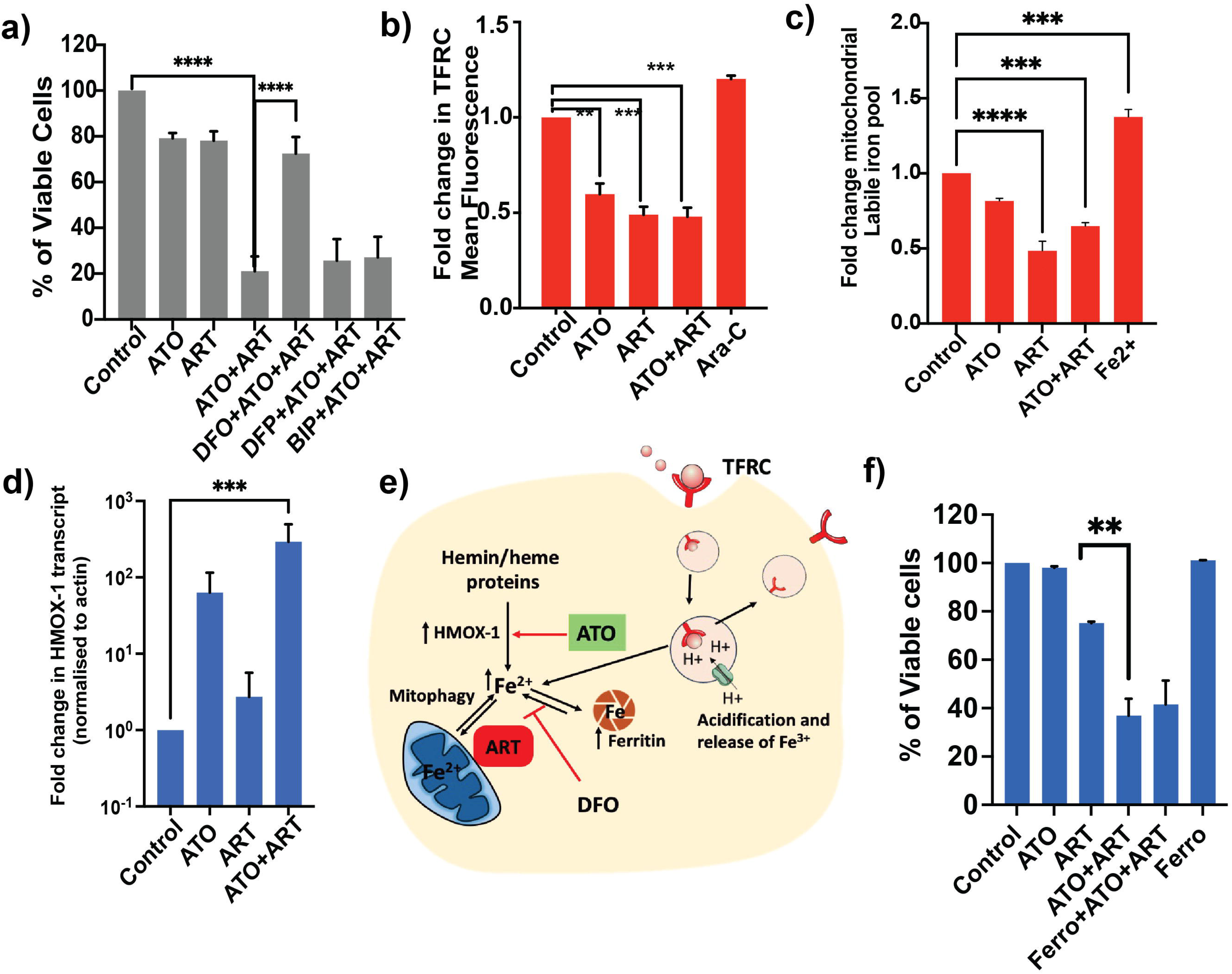
Cellular iron reserves required for anti-leukemic activity of the ART in combination with ATO: a) Viability of U937 cells treated with ATO, ART and iron chelators for 48h (n=4; ATO=2uM; ART=5uM; DFO, DFP and BIP=2uM) b) Fold change in transferrin receptor (TFRC) surface expression mean fluorescence intensity on leukemic cells treated with ATO, ART, ATO+ART and cytarabine (Ara-C) for 24h (n=3; ATO=2uM; ART=5uM; ARA-C = 400ng). c) Fold change in mean fluorescence of MitoFerro green -1 dye intensity that explicitly measures the mitochondrial liable pool levels in U937 cells treated with ATO, ART and ATO+ART, Fe^2+^ as positive control (n=3; ATO=2uM; ART=5uM) d) Transcript levels of haem oxygenase -1 on U937 cells treated with ATO and ART for 24h (n=6; ATO=2uM; ART=5uM) e) Schematic showing the labile iron pool in leukemic cells and effect of ATO and ART. TFRC iron uptake and lysosomal release of iron, Heme based iron supplementation and release of Fe3+ by degradation of Ferritin contributes to enhanced ART activation. Mitochondrial labile iron pool was diminished by ART. Hemin/Heme containing degrading enzyme HMOX-1 is increased by ATO, thereby increasing iron availability for ART activity. DFO – an iron chelator reduced the availability of iron availability for ART activity. (Created with Adobe Illustrator). f) Viability of U937 cells treated with ferroptosis inhibitor (ferrostatin1 – Ferro 1) alone and in combination with ATO and ART for 48h (n=3; ATO =1uM; ART =5uM; Ferrostatin -1 (ferro = 10uM)). Data are presented as mean ± SEM. n.s., P > .05; ∗P < .05; ∗∗∗P < .001; ∗∗∗P < .0001, with a two-tailed unpaired t-test or one-way analysis of variance.

DFO is transported into the cell via an endocytosis process similar to the iron-loaded transferrin receptor. We then measured cell surface expression of transferrin receptor (TFRC) as TFRC-mediated iron uptake is the major cellular iron uptake pathway. Transferrin-bound iron (holo-transferrin) is taken up via the receptor-mediated endocytosis pathway, and endosomes fused with lysosomes, releasing iron in the cytosol and recycling receptors for apo-transferrin to the cell surface. We assessed the effect of ATO + ART on the surface expression of TFRC and labile iron in different iron compartments. ATO and ART as single agents reduced the surface expression of TFRC (figure 4b), indirectly modulating the intracellular iron pool. This effect was only observed with ATO and ART and not with other chemotherapeutic agents. DFO rescued ATO-induced cell death in NB4 cells that are sensitive to ATO but did not impact cell death induced by daunorubicin or cytosine arabinoside (Ara-C) (Figure S3).

Notably, the mitochondrial labile pool (Mito ferro Green-1 – mitochondria-specific labile iron probe) was reduced with ART but not with ATO (figure 4c). In contrast the cytosolic labile iron pool (reported using calcein-AM) was not significantly affected by ATO or ART (data not shown).

We also measured the levels of haem oxygenase 1 (HMOX-1; a metabolic enzyme degrading haem-containing proteins and releasing free iron, bilirubin, and carbon monoxide) which has been shown to be increased by ATO in cancer cells^24^. As expected, treatment of U937 cells with ATO, and ATO+ART increased the HMOX-1 expression (figure 4d). This corroborates the importance of iron for the activity of the combination of ATO+ART. Heme form of iron increases the sensitivity to ART, and ATO enhances this by increasing the expression of HMOX1, while DFO by chelating lysosomal iron rescues the cell from the effect of this drug. The efficacy of this combination and the pleotropic synergistic mechanism of action of ATO and ART is illustrated in the figure 4e.

We excluded the possibility of ferroptosis as a mode of cell death promoted by ATO+ART by applying ferrostatin 1 (inhibitor of ferroptosis). The combination of Ferrostatin- 1 did not affect the cell death elicited by ATO+ART (figure 4f)

### Iron enhances the anti-leukemic properties of ART

To characterize in detail how iron enhances the activity of ART, we treated U937 cells with transferrin (holo and apo forms), Iron II sucrose, hemin (protoporphyrin IX containing Fe^3+^) and δ-aminolaevulinic acid (ALA). We noted that only heme-based iron supplementation (using hemin and ALA which are predominantly in the mitochondria) showed synergy in enhancing the apoptotic activities of ART and ATO+ART (figure 5a, S4a and S4b). We observed similar effects on primary bone marrow specimens of AML (figure 5b). Interestingly, the presence of ART led to low levels of the iron storage protein ferritin (FTH), while ATO increased the FTH thereby corroborating the importance of iron in their synergism (figure 5c). Our, published TMT-labelled proteomics of primary leukaemia samples compared to normal peripheral blood cells showed that the leukemic cells had increased iron-related proteins (FTH, FTL, TFRC, FECH and TOM20)^25^ (fig 5d). To assess clinical relevance, we interrogated the Cancer genome ATLAS data of AML and noted that higher expression of FTH at diagnosis poorer the overall survival (figure 5e). The endoperoxide moiety of the artemisinin family plays a vital role in the antimalarial and anti-leukemic activity of ART and its active metabolite dihydroartemisinin (DHA). The endoperoxide moiety reacts with iron and heme, resulting in an iron-deprived environment, as evidenced by the accumulation of protoporphyrin IX (PpIX – a fluorescent metabolite)^26,27^. To confirm this, we tested a deoxy derivative of artemisinin lacking the reactive endoperoxide ring with iron, and it was inefficient in eliciting the fluorescence that was observed with ART and the active metabolite dihydroartemisinin (figure 5f). These observations are summarized the illustration Figure 5g.

**Figure 5:**
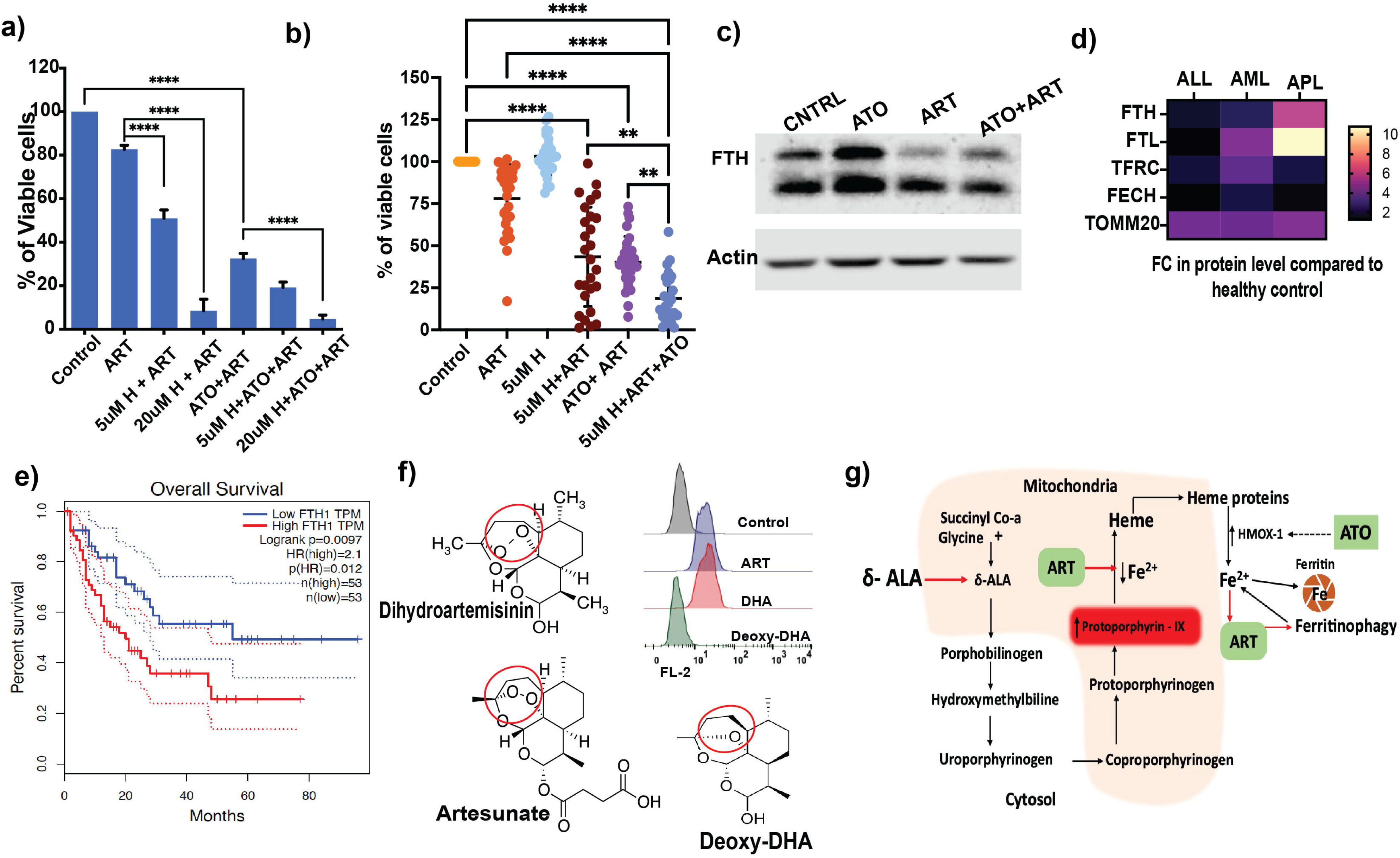
The specificity of the combination therapy is shaped by the decisive presence of intracellular iron. a) Viability of U937 cells treated haemin in combination with ART and ATO+ART for 48h (n=4; ATO = 2uM; ART=5uM; Haemin =5uM and 20uM). b) Viability of primary AML cells treated with haemin, ART, ATO, and combinations for 48h (n=27; ATO = 2uM; ART=5uM; Haemin =5uM) c) Immunoblot of FTH in U937 cells post-exposure to ATO, ART and ATO+ART for 24h (n=3; ATO =2uM; ART = 5uM). d) Heatmap of TMT-labelled proteome (FTH, FTL, TFRC, FECH, TOMM20) of primary AML, APL and ALL cells compared to healthy control mononuclear cells. (5 samples per group, and the differentially expressed proteins and fold change were calculated compared to healthy control mononuclear cells). e) Kaplan-Meier curves of overall survival in TCGA AML patients in the FTH-high red (red) and FTH-low (blue) groups. f) Structure highlighting the endoperoxide moiety of artesunate (ART), active metabolite dihydroartemisinin (DHA) and the inactive metabolite deoxy dihydroartemisinin (Deoxy DHA), histograms illustrating the U937 cells treated with these three agents for 24h and the fluorescent intermediate acquired in flow cytometry. (n=3; 5uM= ART, DHA, and Deoxy derivative) g) Protoporphyrin –IX (PPIX) is a fluorescent intermediate of the heme biosynthesis pathway. The precursor of heme biosynthesis a-ALA enhanced ART activity as seen with heme based supplementation. Due to ART utilisation of iron that is required for heme result in accumulation of fluorescent intermediate PPIX that is observed with ART and its active intermediate DHA not with the deoxy form ART/DHA. Hemin/Heme containing degrading enzyme HMOX-1 and ferritin is increased by ATO thereby increasing the availability of iron for ART activity (Created with Adobe illustrator) Data are presented as mean ± SEM. n.s., P > .05; ∗P < .05; ∗∗∗P < .001; ∗∗∗P < .0001, with a two-tailed unpaired t-test or one-way analysis of variance.

### Novel ART-based combinations are effective in AML

Azacytidine (AZA) combined with venetoclax, is used as front-line treatment for elderly patients and those with a poor performance score who are considered unfit to receive a standard intensive 7/3 induction therapy^28^. We evaluated ATO+ART with AZA to assess the potential role of this clinically relevant combination on primary AML samples, AML and ALL cell lines and confirmed that these drug combinations are not antagonistic (Figure 6a). ATO+ART+AZA had minimal effect on the normal peripheral blood mononuclear cells (figure S5). We evaluated the in vivo efficacy of ATO+ART and ATO+ART+AZA in a THP-1 AML cell line-derived luciferase xenograft model (figure 6b). Tumor burden was monitored using bioluminescence imaging every 3 days of therapy. There was a significant reduction in overall tumor burden with ATO+ART and an even more pronounced effect with ATO+ART+AZA (figure 6c and 6d).

**Figure 6:**
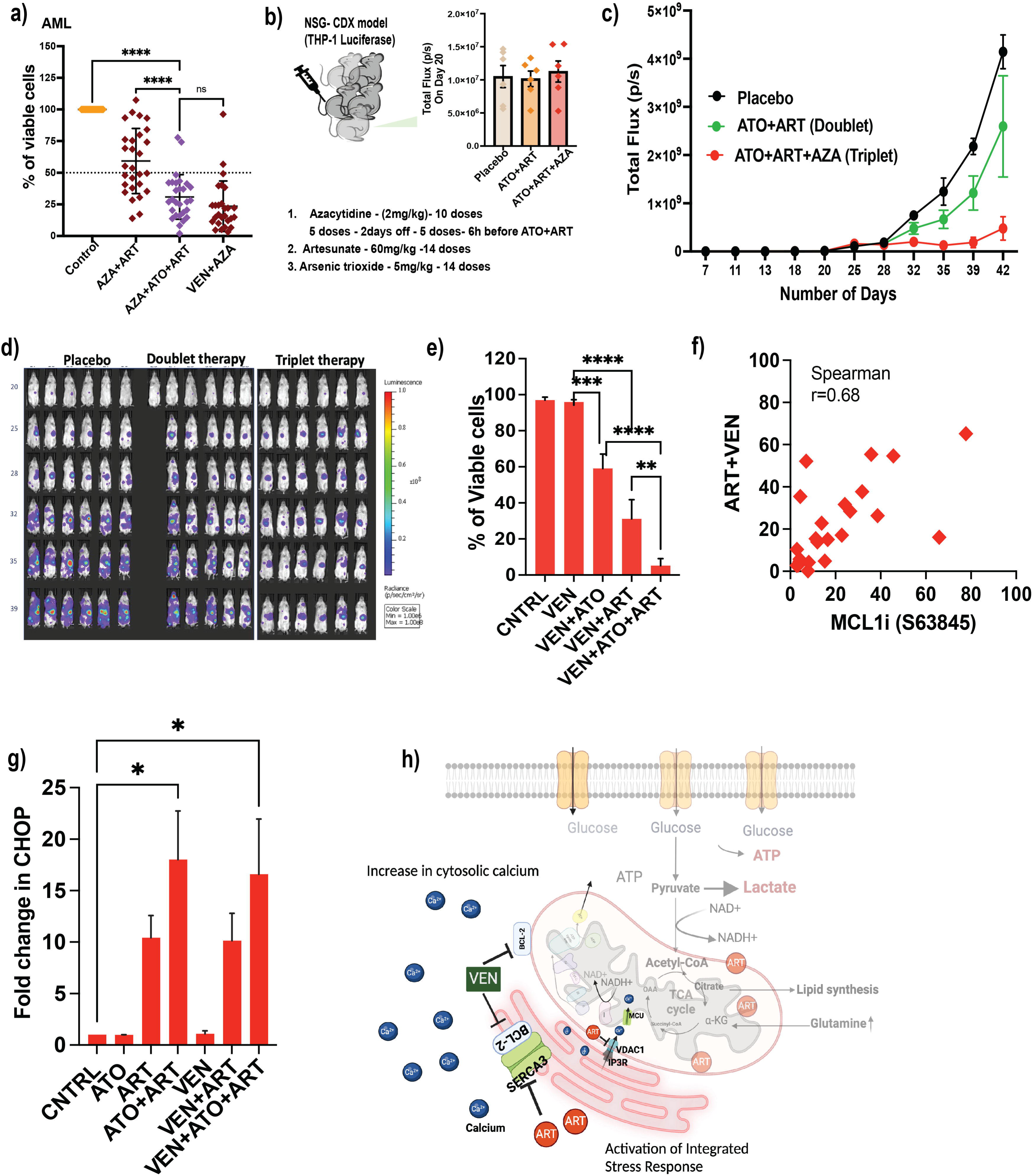
Novel ART-based combinations are effective in AML: a) Viability of primary BM AML samples treated with VEN, ART, ATO, AZA, and rationale combinations for 48hrs (n=28; ATO=2uM; ART=5uM; VEN=500nM; AZA = 2.5uM). b) THP-1 luciferase cell line-derived xenograft model and the leukemic burden before initiation of combination therapies (6 animals per experimental group). c) Bioluminescent intensity of photons emitted from each mouse in the images was quantified during the experiment. Tumour growth was monitored every 4 days by bioluminescence imaging in THP-1 luciferase-engrafted mice and the experimental groups (Black – placebo; Green – ATO+ART and Red – ATO+ART+AZA). d) Bioluminescent images of mice transplanted with THP-1 luciferase cells. Mice were administrated with vehicle, ART+ATO and ATO+ART+AZA Venetoclax (Ven) from day 21 to 34 post transplantation as described in figure 6b. The same mice are depicted at each time-point (n=6 mice per group). e) Viability of U937 cells treated with VEN, ART and ATO for 48h (n=3; ATO=2uM; ART=5uM and VEN= 500nM). f) Spearman correlation (right) of cell death observed in primary AML specimens treated with ART+VEN versus MCLi inhibitor (n=23; ATO=2uM; ART=5uM; VEN=500nM; MCLi= 500nM) for 48h. g) Transcript level of CHOP – ER stress response gene post 24h treatment with ART in combination with ATO, VEN and ATO+ART+VEN (n=3; ATO=2uM; ART=5uM; VEN=250nM) h) **Hypothetical model depicting the synergy of ART and VEN:** ART’s known anti-malarial target SERCA3 – ER Ca^2+^ import channel is negatively regulated by the direct interaction of the anti-apoptotic protein BCL-2. In VEN resistant MCL-1 dependent cells (U937 and primary AML cells), ART in combination with VEN promoted cell death. Cell death induced by this combination could be due to dysregulated Ca^2+^ distribution and the activation of integrated stress response. Distribution of Ca^2+^ is tightly regulated and ART effect on leukemic cell Ca^2+^ dysregulation and its synergy with VEN warrants further studies. (Created with biorender.com) Data are presented as mean ± SEM. n.s., P > .05; ∗P < .05; ∗∗∗P < .001; ∗∗∗P < .0001, with a two-tailed unpaired t-test or one-way analysis of variance.

We also evaluated the effect of ART in combination with venetoclax (VEN, BCL-2 inhibitor) because recent studies have established that inhibition of SERCA3 overcomes VEN resistance by interfering with Ca^2+^ signaling pathways^29^. Additionally, a specific inhibitory effect of ART on SERCA3 activity has been shown in different cellular models, consistent with its original proposed antimalarial mechanism of action as a SERCA(PfATP6) inhibitor. As expected, ART synergized well in combination with VEN in promoting apoptosis even in a VEN- resistant AML cell line (U937) and matched primary AML samples (Figures 6e and 6f). The effect of VEN+ART or VEN+ATO on primary human AML specimens was observed to be equivalent to the effect seen with MCLi inhibition and the magnitude of the cell deaths was highly correlated (figure 6g; Spearman r=0.68). ART induced energy crisis activated integrated stress response (ISR) and synergized with ATO and VEN in promoting cell death of the leukemic cells. It was confirmed by the upregulation of ISR genes ATF4 and CHOP by ART and its combination with ATO and VEN post 24hrs of treatment on U937 and NB4 cells (Figure 6h, and i). ART impacts multiple aspects of AML CEM, including mitochondria, iron metabolism, and SERCA3; additionally, it is relatively specific to cancer cells in comparison to normal counterparts. These observations make ART an attractive, and safe candidate to exploit for anti-leukemic combination therapies with ATO, VEN and or AZA.

Our findings suggest the final common pathway for these combination therapies ART based novel combination is through integrated stress response and subsequent derangement of mitochondrial calcium homeostasis (Figure 6j). VEN and ART act on co-localized proteins (BCL2 and SERCA3, respectively), whereas ATO acts in BCL2-dependent and independent pathways^30,31^.

## Discussion

Affordable and safe treatments for AML and those that can overcome resistance to conventional treatment regimens are urgently needed. We have implemented a drug repurposing strategy that can readily be adapted for clinical trials to assess potential benefits for leukemias as rapidly as possible. ART is a very well-tolerated and safe antimalarial that is being developed for additional antiparasitic, antiviral, and other anticancer indications. ART has many targets in different systems, including SERCAs in malarial and mammalian cells^23^. ART also targets mitochondria, enhances mitophagy^32^, promotes cytochrome-c release^33^, and interferes with many master regulatory pathways in cancer models ^34^.

ART, like FCCP, dissipates mitochondrial membrane potential (MMP)^13^, so we validated it as a mitochondrial uncoupler, selectively targeting leukemic cells. AML cells may escape reliance on OXPHOS by enhancing glycolysis. Inhibiting glycolysis should increase the killing of ALL, AML and APL cells by ART, as confirmed using ATO, which inhibits several glycolytic enzymes, including the rate-limiting step of pyruvate dehydrogenase^35,36^.

Chemical drug proteomics revealed that ART has pleiotropic targets involved in fatty acid synthesis, degradation, and mitochondrial pathways. These findings are consistent with previous studies in solid tumors and malarial parasites^37–39^. Mitochondrial proteins such as CLPB and VLCAD ^18,40^, which are potential therapeutic targets in AML cells, were also labeled by ART. Genetic inhibition of VLCAD has been shown to increase PDH activity in AML cells, single agent ATO-induced cell death in the VLCAD knockdown AML cells (Fig 3f) corroborates this observation,

ART treatment disrupted mitochondrial cristae in AML cells and promoted mitophagy. Mitophagy is essential to maintain the mitochondrial structure on which MMP depends to drive ATP production. Consistent with this in our experiments when ART was combined with a mitochondrial fission inhibitor (mdivi-1) to inhibit mitophagic clearance there was increased cell death. Similarly, cellular pools of NAD^+^/NADH regulate OXPHOS, and an increased NAD^+^ pool is associated with relapsed AML^41^. Availability of NAD^+^ is critical for NAD-dependent enzymes of the TCA cycle and glycolysis^42^. Supplementation of NAD^+^ partially reduced the anti-leukemic efficacy of ATO+ART (Fig 3f).

Intracellular iron activates ART and increases antimalarial and anti-cancer efficacy^43^. Increased iron uptake via transferrin (TFRC) delivered to intracellular iron maintains cell proliferation and respiration in AML cells. DFO abolished the anti-leukemic activity of the ATO+ART (Fig 4a), unlike BIP and DFP, which target non-lysosomal compartments and have lower affinities for iron. ATO enhanced iron availability for ART activation by increasing levels of HMOX-1 and inhibited glycolysis to promote cell death. ART preferentially quenched and reduced the mitochondrial labile iron pool and promoted cell death independent of lipid peroxidation associated with ferroptosis. We studied this mechanism in greater detail because others have shown that ART promotes ferroptosis. DeoxyDHA is not activated by iron and has no antimalarial or anticancer efficacy, even in combination with ATO. Supplementation of iron in the form of δ-aminolevulinic acid or haemin enhances the anti-leukemic properties of ART even without ATO. ART also quenches labile iron, which results in the accumulation of fluorescent intermediates of heme biosynthesis detectable by flow cytometry. As expected, deoxyDHA did not increase fluorescent intermediates for heme synthesis, and DFO successfully abrogated this accumulation observed with ART (Fig 5f). We and others have shown that AML cells have increased intracellular ferritin and mitochondrial respiration ^44^. Increased FTH expression in TCGA AML patients was also associated with poor prognosis.

VEN+AZA is the favoured induction therapy for elderly patients who cannot tolerate 7+3 induction ^3^. ART+ VEN enhance cell death even in VEN-resistant cell lines. Similar efficacy was observed with ATO+VEN in primary AML cells, and both combinations are correlated, albeit less strongly, with killing by a specific MCL-1 inhibitor (S63845). VEN specifically targets BCL-2, an anti-apoptotic protein crucial for the survival of AML cells^45,46^.BCL-2 has been co-localised with SERCAs and has recently been shown to have crucial functional interaction with SERCA3, which is overexpressed in AML exclusively in patients resistant to venetoclax^29^. Interestingly, ratios of SERCA3/SERCA2 are increased in some hematological malignancies, and higher SERCA3 expression portends a poorer prognosis (figure S6). SERCA3 is also exquisitely sensitive to inhibition by ART when expressed in HEK2 cells^47,48^. These observations suggest the following model: ART can act as a VEN mimic by inhibiting SERCA3 in AML cells. We suggest both VEN and ART exert their antileukemic effects by disrupting calcium homeostasis and, thereby, mitochondrial respiration, and combining either or both drugs with ATO has similar and potentially clinically useful efficacy, even when there is VEN resistance.

There is also a rationale for combining ATO, VEN, and ART to achieve maximum therapeutic effect, even in VEN resistant AML while minimize the risk of off-target adverse events. Clinical trials are now planned to assess some of these novel combinations.

## Acknowledgements

Intas Pharmaceutical (India) and NATCO Pharmaceutical (India) for providing active pharmaceutical ingredients of pharmaceutical drugs for this study.

Prof. Tamil Selvan, PhD, Department of Biotechnology, Anna University (Chennai, India), for providing access to the seahorse extracellular flux analyser.

Central Imaging Facility and Flowcytometry Facility, National Centre for Biological Sciences and Dr.Ganesh Kadasoor from Olympus for the confocal imaging experiments.

Central Electron Microscopy facility, Christian Medical College, Vellore, Tamil Nadu, India. IA/CPHS/18/1/503930 – Funding granted to VM from India Alliance Wellcome DBT (Department of Biotechnology), New Delhi, India.

BT/IN/UK/DBT-BC/2017-2018- Funding granted to NB by Newton PhD fund governed by British Council, United Kingdom.

The Seahorse facility at Anna University is funded by the Department of Science and Technology Fund for Improvement of Science and Technology (S&T) Infrastructure (SR/FST/LSI-649/2015).

## Author contributions

N.B, S.K and V.M conceived this work. N.B, with assistance from A.V and H.K performed the cellular, molecular, proteomics, primary experiments, and metabolic assays. C.R performed the metabolomics and data analysis. Original Manuscript draft: N.B, S.K and V.M. Writing – Reviewing and editing of the manuscript: N.B, Y.A, S.K and V.M. Funding Acquisition: N.B, S.K and V.M. Resources are provided by A.K, U.K, P.B, E.S, B.G, A.A, S.K and V.M. The study was supervised by S.K, V.M and Y.A.S.K. and V.M. equally contributed to this work.

The project is a highly interdisciplinary initiative to repurpose artesunate as a treatment option for acute myeloid leukemia. This work has been carried out in both V.M’s and S.K’s laboratories and reflects their respective expertise in leukemias and drug repurposing with artesunate and therefore their co-corresponding authorships.

## Data and code availability

All data reported in this paper will be shared by the lead contact upon request. This paper does not report original codes.

## Lead contact

Further information and requests for resources and reagents should be directed to and will be fulfilled by the Lead Contact, Dr. Vikram Mathews.

## Supplemental information titles and legends

**Document S1:** Figure S1-S6 and methods.

**Table S1:** Relative protein abundances enriched in the biotinylated ART pull-down of U937 cells (related to figure 2b and c).

**Table S3:** Metabolites peak intensities of ART-treated U937 and control cells (related to supplementary figure 2).

## Methods

### Cell culture experiments

All cells were grown at 37°C in a 5% CO2 humid atmosphere. NB4 (a kind gift from Dr. Harry Iland, RPAH, Sydney, Australia, with permission from Dr. Michel Lanotte), NB4 EV-AsR1, NB4 EV-AsR2 (in-house generated ATO resistant cell line), MV-411 (ATCC), THP-1 (ATCC), U937 (ATCC), and Jurkat (ATCC) cells were cultured in RPMI-1640 medium (Sigma) containing 2 mM L-glutamine, supplemented with 10% fetal bovine serum (ThermoFisher Scientific) and 1× penicillin-streptomycin (ThermoFisher Scientific). Kasumi-1 (ATCC) cells were cultured in RPMI-1640 medium containing 2 mM L-glutamine, supplemented with 20% fetal bovine serum and 1× penicillin-streptomycin. SUP B15 (ATCC) cells were cultured in Iscove’s Modified Dulbecco’s culture medium (Sigma) supplemented with 20% fetal bovine serum and 1× penicillin-streptomycin (ThermoFisher Scientific). Cell lines were regularly passaged every 2-3 days for maintenance in the exponential growth phase and tested for mycoplasma contamination every 6 months in the laboratory using a Universal Mycoplasma detection kit (ATCC).

### Primary human Specimens

Bone marrow samples were collected from patients diagnosed as AML, ALL, and APL at Christian Medical College who had been newly diagnosed. Samples were collected after informed consent was obtained, and studies were conducted under the protocols approved by the Institutional Research Ethics Committee (IRB Min No -11796). Mononuclear cells were isolated using Ficoll density gradient. Freshly processed or thawed cryopreserved cells were used in drug sensitivity assays. The primary samples were cultured in Iscove’s modified Dulbecco’s culture medium (Sigma) supplemented with 20% fetal bovine serum and 1× penicillin-streptomycin (ThermoFisher Scientific).

### Animal Studies

NSG (NOD.Cg-PrkdcscidII2rgtm1wjl/SzJ) were purchased from Charles River Laboratories. 1x10^6^ THp-1 luciferase viable cells in 100µL PBS were administered by intravenous injection via the tail vein. Both male and female mice aged 10-12 weeks of old were used in this study. Tumour growth was checked twice weekly (from day 7) by bioluminescent imaging (BLI). Briefly, the mice were injected (s.c) with 150 mg/kg d-Luciferin 15 minutes prior to imaging. 10 minutes following administration of d-Luciferin mice were anaesthetized and placed into the imaging chamber (Spectrum CT) and imaged for luminescence. The imgaes were captured and processed using Living Image software (Caliper LS, US). Mice were randomly allocated to treatment groups. The treatment commenced once progressive tumour growth was evident (Day 20), and randomization was based on bioluminescence. The drugs were administered intraperitonially, and the treatment continued for two weeks with an 11-day observational phase followed by a further 5 days of dosing. The Crown Biosciences United Kingdom performed the experiments according to their CBUK guidelines and ethical approval.

### Cell death assay

5 × 10^5^ cells were treated with different drugs and inhibitors and incubated for 48 hours at 37 °C CO2 incubator. After the incubation period, the cells were collected, and the viability was assessed using Annexin V/7-amino actinomycin D (7AAD) apoptosis assay kit (BD Pharmingen) as per the manufacturer’s protocol.

### Seahorse metabolic flux analysis

Extracellular flux assay kits XF24 (Agilent Technologies, CA, USA) were used to measure oxygen consumption and glycolytic flux. 5x10^5^ cells were plated in an XF24 cell culture microplate. Oxygen consumption and glycolytic flux were measured according to the manufacturer’s protocol and as previously described^49^. Briefly, three replicate wells of 5 × 10^5^ cells per well were seeded in a 24-well XF24 plate. At 30 min prior to analysis, the medium was replaced with Seahorse XF Mito stress media (Agilent Technologies) and the plate was incubated at 37 °C non-co2 and performed mito stress test. Analyses were performed at basal conditions and after injection of oligomycin, FCCP, antimycin A, and rotenone.

### Chemical-drug proteomics approach for target identification

Biotinylated-artesunate (B- ART) was synthesized by linking PEG-Biotin to the dihydroartemisinin (active metabolite of artesunate) and used to pull down the interacting proteins and streptavidin on bead digestion were carried out. U937 cells were treated with B-ART for 6 hours in combination with ATO at the end of incubation the cells were lysed using MS compatible lysis buffer (50mM Tris (pH 8.0), 150mM NaCl, Protease inhibitors) and clear supernatant were collected and proceeded with streptavidin pull down to enrich biotinylated proteins as per the manufacturers protocol (#Thermofisher scientific). On column trypsin digestion were performed on the samples. Non-biotinylated control samples were used as a control to identify and exclude non-specific targets of biotinylated ART. Peptides were identified and quantified using protein pilot software 4.5(SCIEX) with a paragon algorithm in reference to the International Protein Index (IPI database).

### Immunoblot

Cell homogenates were obtained by cell lysis in IP Lysis buffer (Thermo Scientific), with complete protease inhibitors (Roche, Basel, Switzerland). Protein was quantified using Bradford assay (Bio-Rad). 25ug of lysates per lane was loaded were analysed in SDS–PAGE. After protein transfer to a nitrocellulose membrane, membranes were blocked with non-fat dry milk (5%, 2 hours) and incubated with primary antibodies overnight. After an overnight incubation the membranes were washed thrice with wash buffer containing tween 20 and incubated with HRP conjugated anti-mouse and anti-rabbit secondary antibody for an hour. The membranes were washed 5 times, and the protein bands were detected by the standard chemiluminescence method (Thermo Scientific).

### Mitochondrial morphology using confocal laser scanning microscopy

MitoTracker™ Red CM-H2Xros (Thermo-Scientific) was added into culture media at a final concentration of 100nM to label the mitochondria of leukemic cells for 30 minutes in 37o C 5%CO2 incubator before harvesting cells for confocal studies. To prepare slides, labeled cells were washed 2 times with ice-cold PBS and re-suspended at a final concentration of 500K cells /ml in ice-cold PBS. At room temperature, 150 µl of cell suspension (75K cells) was used for cytospin at 500rpm for 5 minutes. The slides were air-dried in the dark box at room temperature for 5 minutes before being fixed in -20° C 100% methanol for 10 minutes. The slides were then air-dried in dark box at room temperature for 10 minutes and mounted in Vectashield DAPI mountant before confocal imaging. The stained slides were imaged on the Olympus FV3000 confocal microscope using its high sensitivity detectors excite the DAPI and its 633nm laser. Z stacks were captured, and the images were deconvoluted using Olympus Fluoview.

### Measurement of mitochondrial Labile iron pool

Briefly, 5x10^5^ cells were treated with drugs and after 6 hours, the cells were stained with Mito Ferro green-1 (Dojindo Molecular Technologies, Inc.) for 15 minutes in Phosphate buffered saline (PBS) at 37°C followed by washing with PBS and the fluorescence intensity was measured in FACS Calibur with an excitation wavelength at 485nm, and the data analysed using Kaluza Analysis Software (Beckman Coulter).

### Semi-quantitative real-time polymerase chain reaction

Total RNA was extracted using Trizol reagent (Invitrogen Carlsbad, CA, USA). 500ng of the extracted RNA was converted into cDNA using superscript II cDNA kit (Invitrogen Carlsbad, CA, USA). The expression of genes was studied using SYBR green method (Finnzymes F410L, Thermo scientific, Rockford, IL, USA). The Ct values were normalized with actin and the fold differences were calculated using 2-ΔΔCt method.

### Transmission electron microscopy

Two million leukemic cells were collected post 24hours treatment or control and washed twice with phosphate buffer. The cells were then fixed with a fixative -4 % paraformaldehyde plus 3% glutaraldehyde in 0.1 M phosphate buffer pH 7.2 (PB)-for 30 minutes at room temperature and 4°C overnight. Cells were then washed with phosphate buffer three times for 15 minutes. Next, cells were post-fixed with 1% osmium tetroxide buffered with PB for 1 hour and washed again using PB twice for 20 minutes, dehydrated with an ethanol series, then infiltrated with propylene oxide. The samples were then resin embedded with Epon, which was polymerised at 60°C for 48 hours. Solid epoxy blocks were sectioned on a Leica Ultracut microtome to 100 nm thickness, The ultra-thin sections were collected on copper grids and stained with uranyl acetate and Reynolds solution (sodium citrate and lead citrate) to give contrast. Sections were transmitted in electron microscopy Tecnai T12 spirit and photographed.

### Quantification and statistical analysis

Statistical analyses were performed using GraphPad Prism version 9.3.1. Data are presented as mean ± SEM. All data points are represented as mean ± standard error of the mean. Two-tailed Student t-test was used to compare mean values between the 2 groups. A one-way analysis of variance was used for experiments in which multiple groups were compared with the control group. Results are considered significant when ∗p <0.05, ∗∗p <0.01, ∗∗∗p <0.001, and ∗∗∗∗p <0.0001.

