## Supplementary File 1 for "Targeting Metabolic Dependencies in Acute Myeloid Leukemia: A Dual Strategy using Arsenic Trioxide and Artesunate"

**Supplementary figures and Methods:**

**Supplementary Figure 1:**

a) Representative images of Wright-Giemsa stained U937 cells treated with ATO, ART and ATO+ART for 24h (ATO=2uM; ART=5uM).

(b-c) Reactive species levels of U937 cells treated with ATO, ART and ATO+ART for 6h and 24h (n=3; ATO=2uM; ART=5uM).

d) Viability of U937 cells treated with ATO, ART and ATO+ART in combination with CB839 and etomoxir for 48h (n=6; ATO=2uM; ART=5uM and CB839= 10uM; ETO= 10uM).

(e-g) Relative expression levels of GLUT-1, CD36 and GLS in U937 cells treated with ATO, ART and ATO+ART. (GLUT-1 and CD36 - n=3; GLS n=6; ATO=2uM; ART=5uM).

Data are presented as mean ± SEM. n.s., $P > .05$; $*P < .05$; $***P < .001$; $***P < .0001$, with a two-tailed unpaired t-test or one-way analysis of variance.

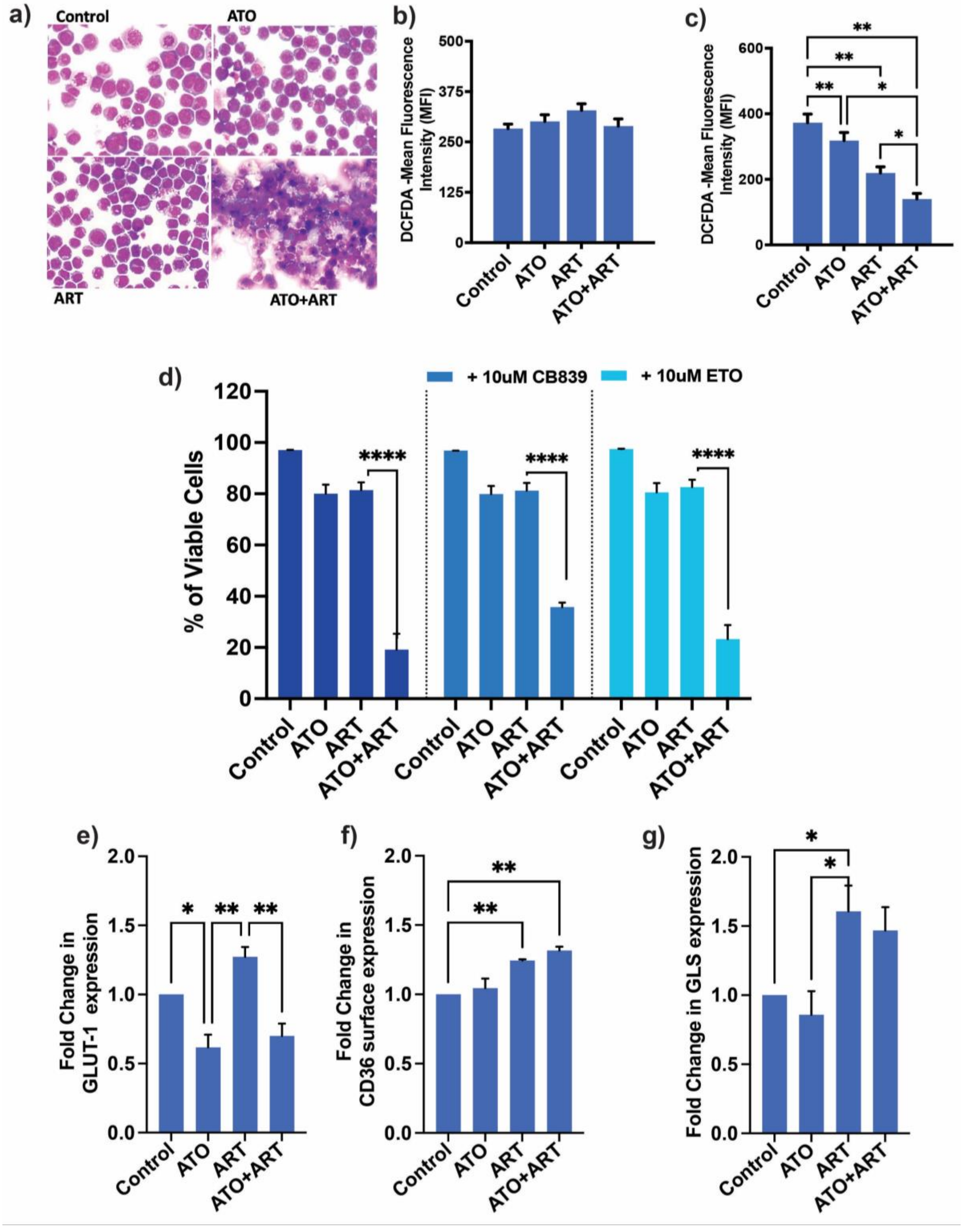
a)
Control
ATO
ART
ATO+ART
b)
DCFDA -Mean Fluorescence Intensity (MFI)
500
375
250
125
0
Control
ATO
ART
ATO+ART
c)
DCFDA -Mean Fluorescence Intensity (MFI)
600
400
200
0
**
**
**
*
*
Control
ATO
ART
ATO+ART
d)
+ 10uM CB839
+ 10uM ETO
% of Viable Cells
120
100
80
60
40
20
0
****
****
****
Control
ATO
ART
ATO+ART
e)
Fold Change in GLUT-1 expression
2.0
1.5
1.0
0.5
0.0
*
**
**
f)
Fold Change in CD36 surface expression
**
**
g)
Fold Change in GLS expression
*
*

**Supplementary figure 2:**

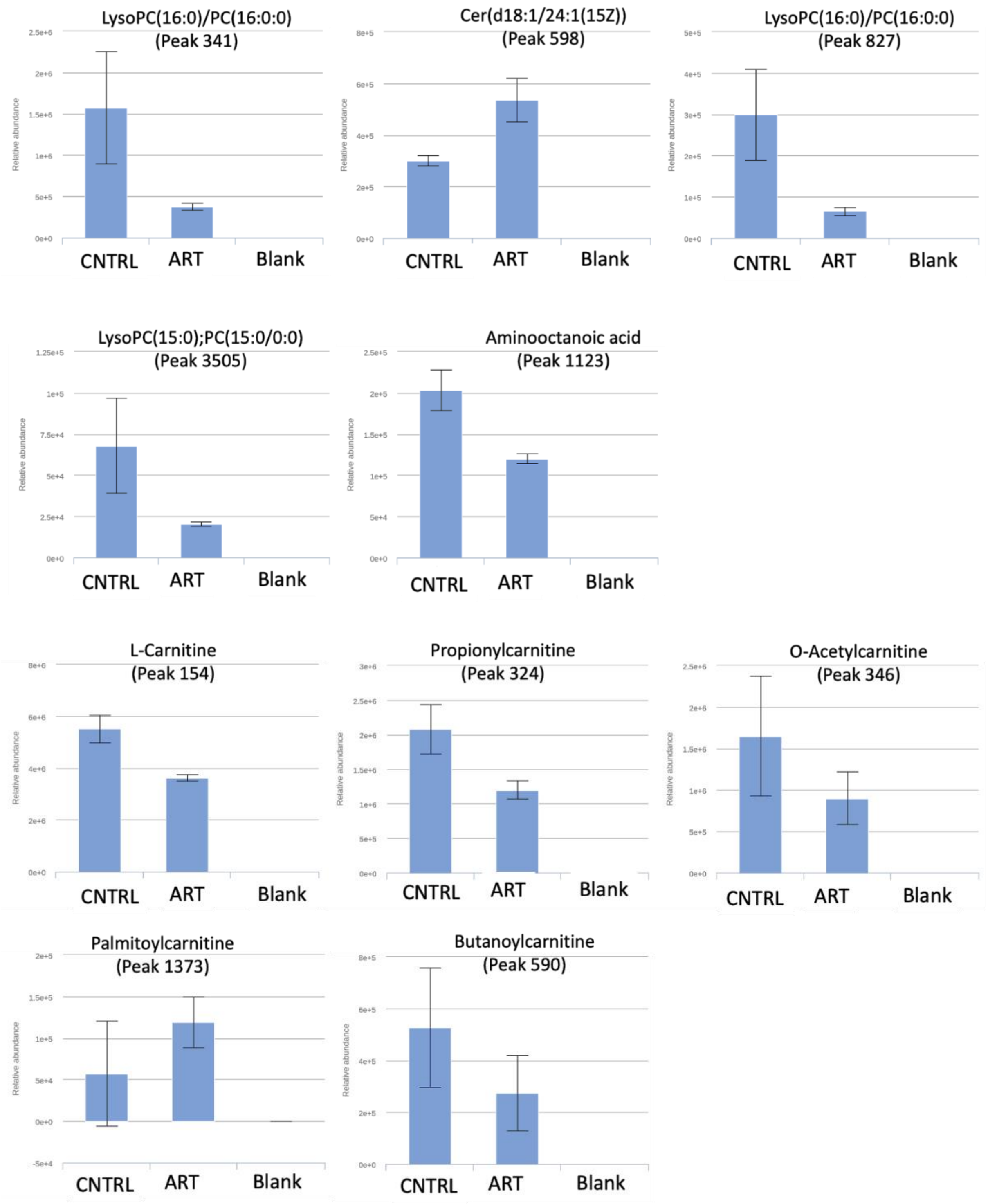

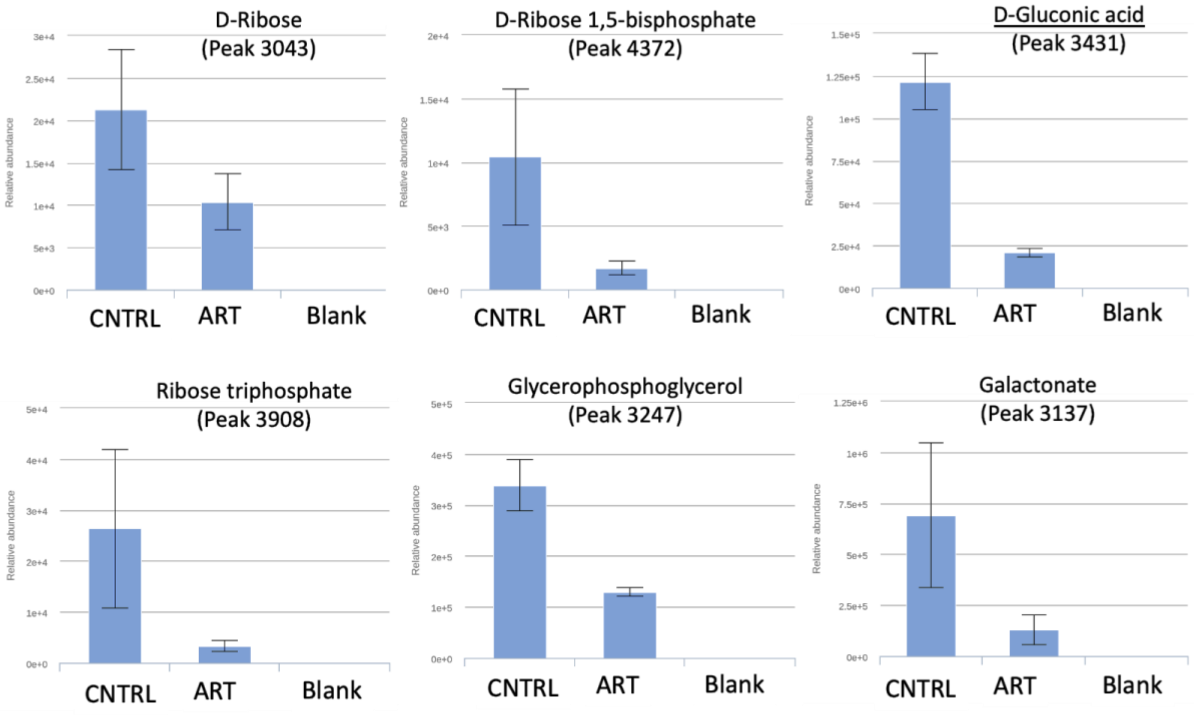


Untargeted metabolomics of U937 cells treated with ART for 24h: Relative abundance of metabolites relating to lysophospholipids, acylcarnitines, and sugars. (n=3; ART=5uM; 24h)

**Supplementary figure 3:**

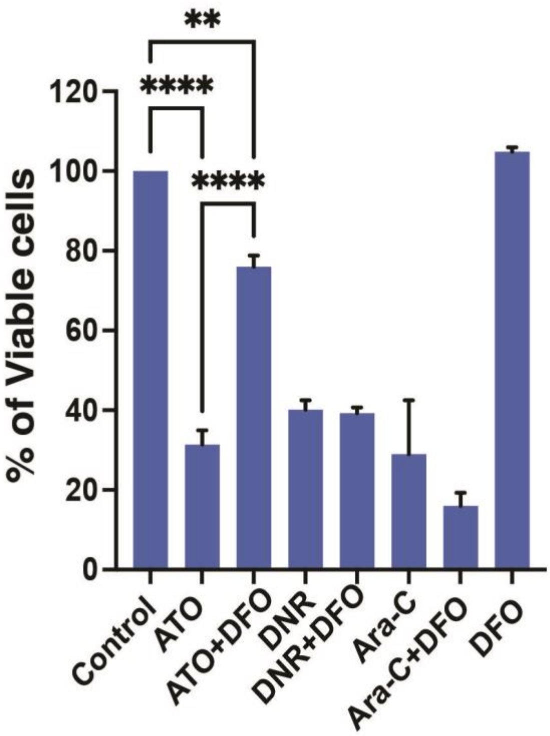


Viability of ATO sensitive NB4 cells treated with ATO, DNR, Ara-C in combination with DFO for 48h (n=3; ATO=2uM; DNR = 40ng; Ara-C =400ng; DFO=20uM).

Data are presented as mean ± SEM. n.s., $P > .05$; ∗$P < .05$; ∗∗∗$P < .001$; ∗∗∗$P < .0001$, with a two-tailed unpaired t-test or one-way analysis of variance.

**Supplementary Figure 4:**

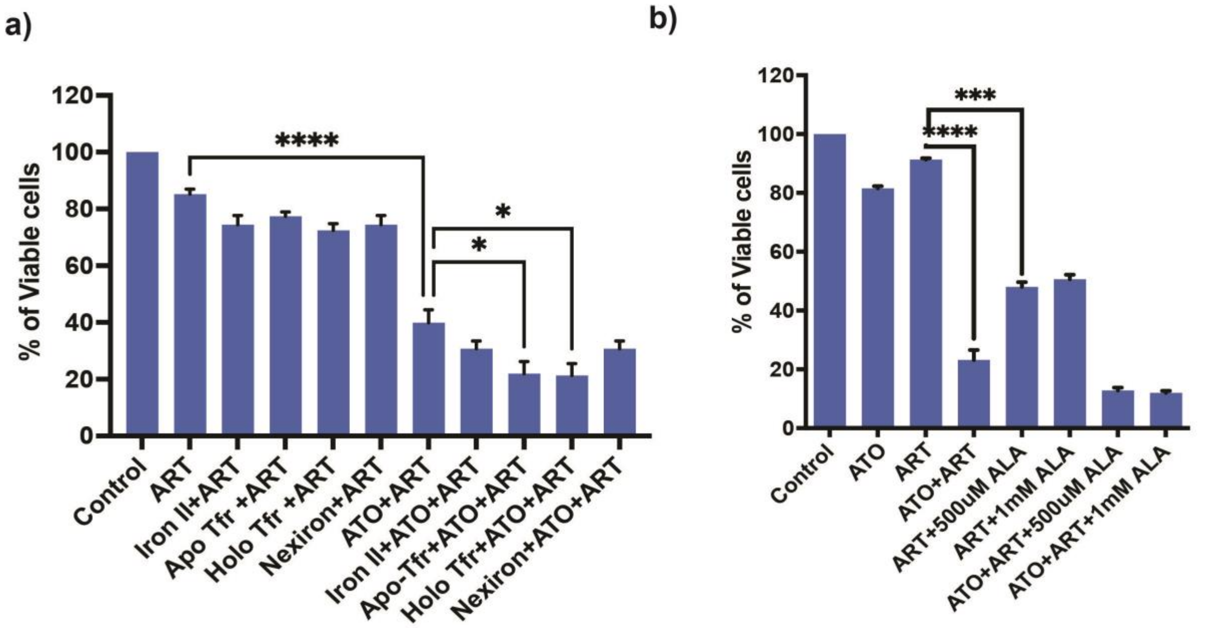


a) Viability of U937 cells treated with different forms of iron in combination with ART and ATO+ART for 48h. (n=3; Iron II = 10ug; Apo and Holo Tfr = 50ug; Nexiron = 40ug).
b) Viability of U937 cells treated with ALA in combination with ATO, ART and ATO+ART for 48h. (n=4; Iron II = 10ug; Apo and Holo Tfr = 50ug; Nexiron = 40ug).

Data are presented as mean ± SEM. n.s., $P > .05$; ∗$P < .05$; ∗∗∗$P < .001$; ∗∗∗$P < .0001$, with a two-tailed unpaired t-test or one-way analysis of variance.

**Supplementary Figure 5:**

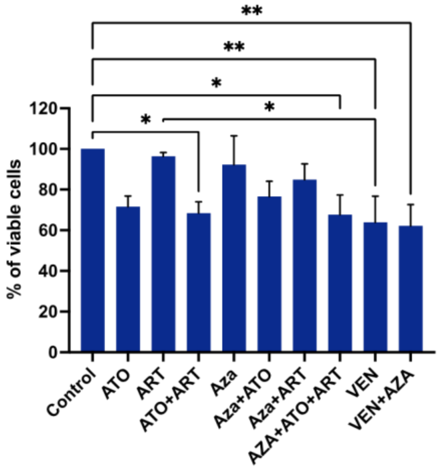


Viability of healthy peripheral blood mononuclear cells treated with ATO, ART, VEN and AZA for 48h. (n=5; ATO=2uM; ART =5uM; Aza =2.5ug; VEN = 500nM).

Data are presented as mean ± SEM. n.s., P > .05; ∗P < .05; ∗∗∗P < .001; ∗∗∗P < .0001, with a two-tailed unpaired t-test or one-way analysis of variance.

**Supplementary figure 6:**

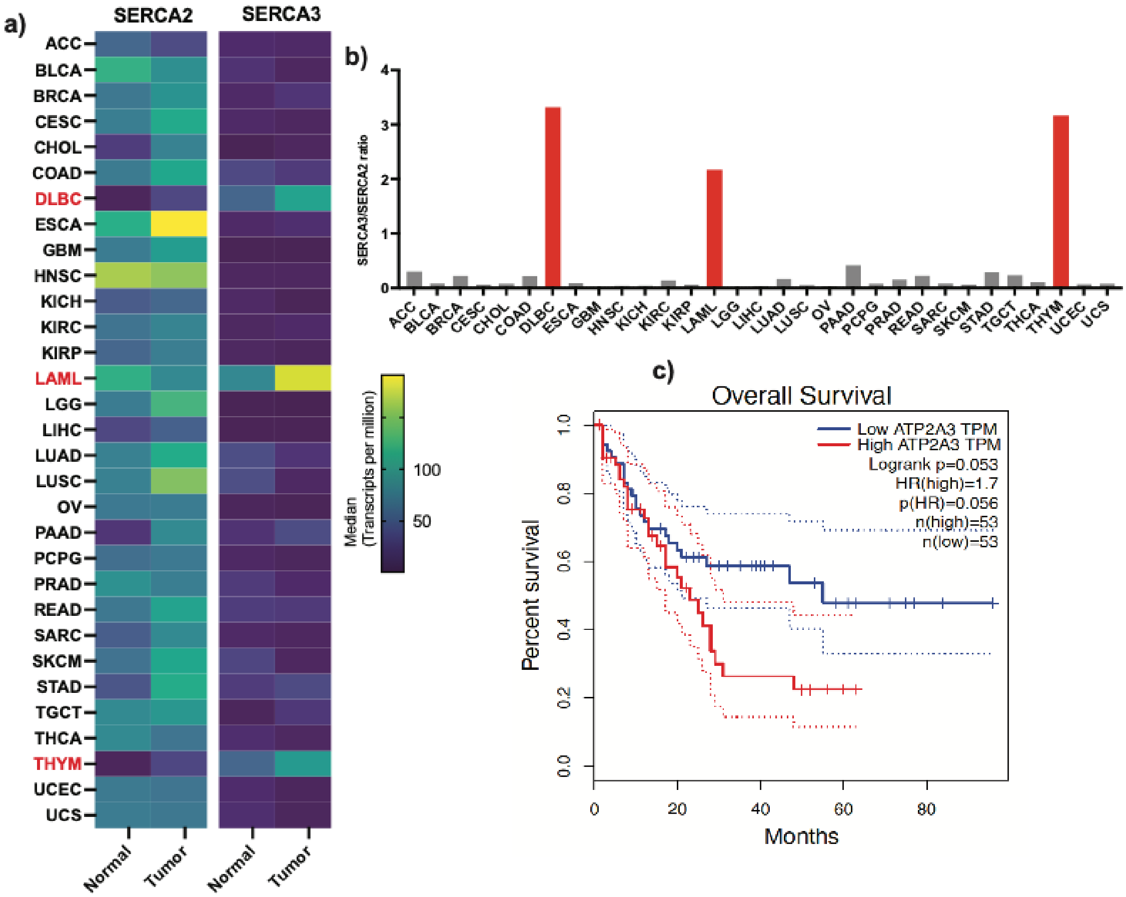


a) SERCA2 and SERCA3 expression levels of normal and cancer tissues from the Cancer Genome atlas TCGA dataset.

b) SERCA3 to SERCA2 ratio for TCGA tumor types. Red bars highlight the hematological malignancies with high SERCA3 to SERCA2 ratio.

c) Kaplan Meier survival curves of TCGA AML cohort stratified based on SERCA3 expression.

**Supplemental Methods:**

**Wright-Geimsa Staining:**

50K U937 cells in one hundred microliters were used to make cytospin slides (500RPM for 5 minutes). The slides were then air-dried and fixed with one hundred per cent methanol for 10 minutes, washed and stained with Wright-Giemsa for 5 minutes, and washed with PBS, and the images were captured in an inverted light microscope using an oil immersion lens (Olympus).

**Endogenous ROS measurement:**

$5x10^5$ treated/control cells were stained with 5uM of DCFDA (Sigma) for 15 minutes at 37 degrees. The stained cells were washed twice with phosphate-buffered saline (PBS) at 500g for 5 minutes. The cell pellet was resuspended in PBS, and the fluorescence was measured using Flow cytometry (Beckman Coulter Navios).

**Untargeted Metabolomics:**

Approximately 5 million cells were cultured and treated with DMSO, ATO, ART and the combination of ATO+ART. After 24 hours of treatment the cells were collected and quenched in a dry ice/ethanol bath followed by centrifugation for 10 minutes at 1000g, 4 degrees. The supernatant was removed completely, and the pellet was re-suspended in 200ul of (1:3:1) ice-cold Chloroform: Methanol: Water mixture to extract the metabolites and incubated for 1 hour in a rocker at 4 degrees. Then, the sample was spun at 13000g for 3 minutes at 4 degrees and the supernatant was collected and stored at -80 degrees. Liquid chromatography-mass spectrometry (LC/MS) for the extracted metabolites was performed using a pHILIC column in a Thermo Orbitrap at the Glasgow polyomics facility, University of Glasgow, UK.
